# Classifying the Unknown: Identification of Insects by Deep Open-set Bayesian Learning

**DOI:** 10.1101/2021.09.15.460492

**Authors:** Sarkhan Badirli, Christine J. Picard, George Mohler, Zeynep Akata, Murat Dundar

## Abstract

Insects represent a large majority of biodiversity on Earth, yet only 20% of the estimated 5.5 million insect species are currently described (1). While describing new species typically requires specific taxonomic expertise to identify morphological characters that distinguish it from other potential species, DNA-based methods have aided in providing additional evidence of separate species (2). Machine learning (ML) is emerging as a potential new approach in identifying new species, given that this analysis may be more sensitive to subtle differences humans may not process. Existing ML algorithms are limited by image repositories that do not include undescribed species. We developed a Bayesian deep learning method for the open-set classification of species. The proposed approach forms a Bayesian hierarchy of species around corresponding genera and uses deep embeddings of images and barcodes together to identify insects at the lowest level of abstraction possible. To demonstrate proof of concept, we used a database of 32,848 insect instances from 1,040 described species split into training and test data. The test data included 243 species not present in the training data. Our results demonstrate that using DNA sequences and images together, insect instances of described species can be classified with 96.66% accuracy while achieving accuracy of 81.39% in identifying genera of insect instances of undescribed species. The proposed deep open-set Bayesian model demonstrates a powerful new approach that can be used for the gargantuan task of identifying new insect species.

---

**D**iversity of life is a central tenet to Biology, from the process of speciation to the maintenance or prevention of extinction (adaptation) and the ecosystem services biodiversity provides. Human activity threatens this, and as a result, the well-being and economics of humans are threatened. Biodiversity is important for health and medicine (3), drug discovery (4), social equality (5), ecosystem services (6), food security (7), and for life (8). The time is now for bold changes to address the current and future losses of biodiversity, but the problem that arises is how to describe the vast amounts of existing biodiversity?

One of the largest groups of animals on the planet is the insects, and they are the most diverse, yet, so few of them are actually described, and they are likely disappearing faster than we can identify them (9). Within Insecta, approximately 5.5 million insects species are thought to exist, yet only 20% are actually described, leaving a very large swatch of unknown biodiversity (1). Describing biodiversity for insects comes down to discovery and identification. Once an insect is collected, an individual with ‘some’ knowledge will try and identify it to the lowest taxonomic level. Higher level morphological characteristics can be quite easy to identify, for example, if it has one pair of wings with the second set of wings reduced to small knob-like structures (halters), one would conclude Diptera, then move on to the next taxonomic level by using published available keys (10). From there, family specific dichotomous keys would allow one to continue until species level is reached, all of which are based on both continuous and discrete characters on the insects. Here is the inherent flaw: undescribed species would not be present in a key, and only through the very thorough analysis of characters could one conclude it may be a new, undescribed species and is not attributed to plasticity or geographic isolation. The use of newer technology, specifically, the DNA Barcode (2), has really helped confirm the new species if the variation in sequence exceeds the traditional intraspecific variation or when species have undistinguishable characteristics such as cryptic species (11).

There are two paths to identifying biodiversity: discovery and identification, and they are interrelated. Advances in technology have addressed one of these to some extent (discovery) through the use of the DNA barcode, however, the species are flagged as unlikely to be an existing species, or they simply exceed the typical intraspecific DNA variation limits, but they have not been identified. One cursory look at the DNA Barcode Database (BOLD) (12, 13)) will reveal that a search of Diptera yields 2.4 million records (DNA sequences) and 126,000 BINs (barcode indexed numbers), yet only 25,000 species have been identified, meaning DNA is facilitating the possibility of new species discovery, but nothing is happening to identify them. So even with DNA sequencing increasing the rate of new species discoveries, we are not identifying them, they are not being published, and the biology around these new species is not being discovered. It just provides a baseline for the approximate biodiversity but does not contribute to the knowledge base.

The increasingly difficult challenge is the lack of experts in a given taxonomic field owing to the vast diversity of the insects themselves, and the art of traditional taxonomy continues to decline, further contributing to the bottleneck (14–16). Therefore, the only way to meaningfully scale the discovery and identification of new species is to address that point, and that is an expert that is carefully trained to recognize and define differentiable characteristics of various insects. If we have the ability to perform this function across a broad scale, the insect identification problem becomes manageable, and this is where machine learning (ML) algorithms can be leveraged to find patterns from insect image databases and apply this to identifying insect species. Although ML can potentially exceed human recognition in exposing subtle morphological features, existing ML paradigms will no doubt be challenged by the scale of the insect identification problem.

Recent advances in ML have led to a surge of interest among scientists in entomology and ML methods have been lately used to provide potential solutions to many challenges in the domain. Deep learning (DL) approaches, in particular involving Convolutional Neural Networks (CNN), are utilized in pest-detection (17, 18), digitalization of Natural History Museum collections (19, 20), measuring invertebrate biodiversity (21, 22), investigating the plant-insect interactions (23) and many more applications (24). There are several studies that explored the use of ML approaches on audio data to classify bee and grasshopper species (25, 26). A recent study also applied CNN on the acoustics of wingbeats to detect the mosquito presence (27). ML methods have also been employed in a more challenging task of automatic detection of species in video and time-lapse images (28). Furthermore, recent studies demonstrated that ML approaches can achieve human-expert level accuracy on image-based taxonomic identification (29–31).

Perhaps the most relevant studies to insect identification and discovery are the ones dealing with open-set recognition. Despite all these great accomplishments, there are very few studies investigating DL methods in an open-set classification setting (32). Furthermore, current open-set methods have been employed on relatively small datasets and do not scale well with a larger number of classes (32). This in turn limits their usefulness in Entomology as insect datasets are very fine-grained and contain a large number of similar classes.

Traditional supervised learning algorithms will be inherently limited by the non-exhaustive nature of insect repositories available for training. It is impractical, often impossible, to create a training repository with a complete set of insect species for various reasons. For example, some of the insect species are not yet described, and thus well-characterized training images of insects from these species simply cannot be obtained. Similarly, when insect species are either rare or non-existent in a given geographical locale collecting samples may become impractical. And finally, insects specifically pose a challenge due to the morphologically distinct life stages of the insect that may include egg, larval and pupal stages.

ML models for insect identification should be designed by recognizing beforehand the non-stationary nature of the biological and environmental processes governing insect emergence, proliferation, and migration. An effective ML model should not only identify future samples of species represented during training (described species) but should also be capable of detecting and identifying samples of unrepresented species (undescribed species). Identifying samples of undescribed species is an ill-defined problem. Although ML models can be tailored to operate in an open-set classification setting to detect insect samples with no matching classes in the training data, such approaches have only been restricted to detect an insect sample as an outlier and cannot differentiate between different types of outliers (33–35).

In this study, we seek to answer whether recent advances in machine learning and computer vision can help extract subtle yet potentially differentiable morphological characteristics, when combined with the highly specific DNA Barcode data, can help facilitate more accurate identification of insects of described species while simultaneously discovering insects of undescribed origin and identify them at the lowest level of abstraction possible (See Fig 1). The core building block of our open-set classification approach is a two-layer hierarchical Bayesian model defined over both described and undescribed species with two different types of priors: global and local. Global prior is shared by all species whereas local priors are only shared by species that are taxonomically more similar and used as a surrogate class for undescribed species. Classification is performed by maximizing posterior predictive likelihood over both true and surrogate classes.

**Fig. 1.**
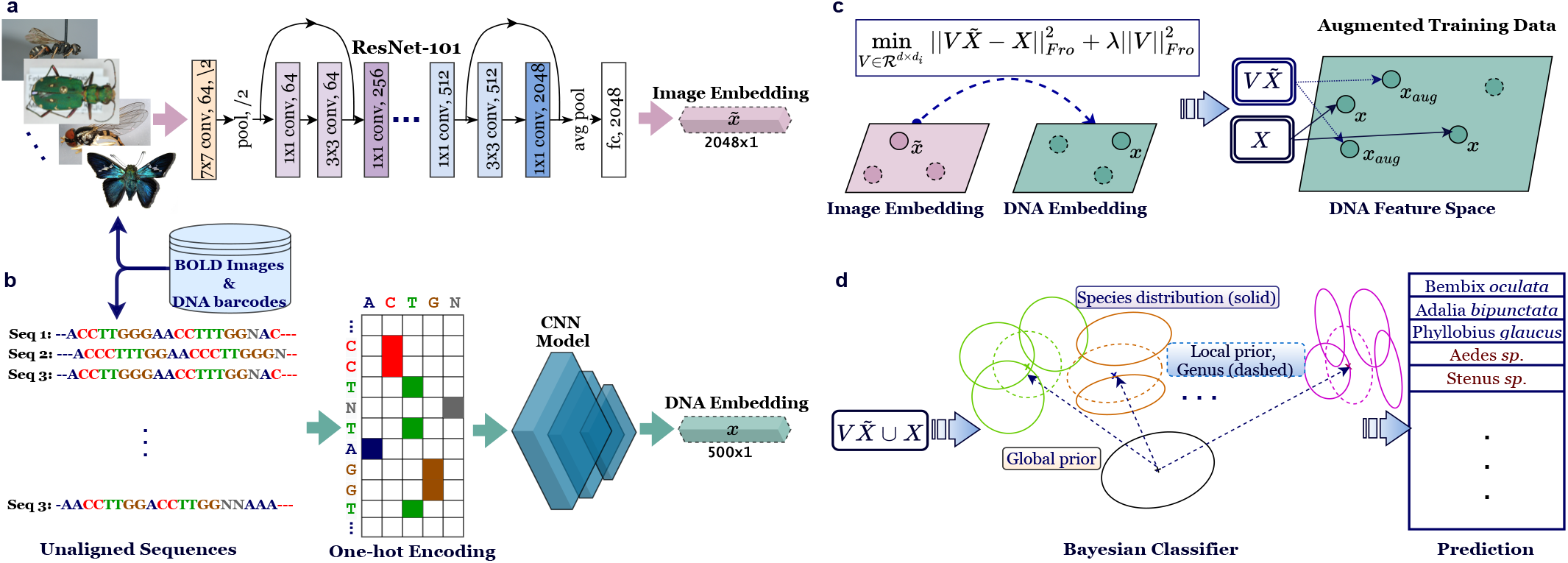
Deep Open-set Bayesian Classification with Unknown and Undescribed Species. **a**. Image embeddings of size 2048 are obtained using the pretrained ResNet-101 model. **b**. CNN architecture is trained using one-hot encoding representations of DNA barcodes (see Supp. Fig 4 for more details). **c**. Mapping from ResNet features to CNN embeddings is learned by transductive Ridge regression. Training set for the CNN embeddings is augmented by the mapped versions of ResNet features. **d**. Open-set Bayesian model is trained on the augmented training set and used for classification. A test sample is either assigned to one of the described species or identified as a new species belonging to one of the described genera.

## Results

In this section, we first briefly discuss the predictive performance of the Convolutional Neural Network (CNN) model we developed to learn DNA embeddings. Then, open-set insect classification results are reported and, finally, the section is concluded with discussion and case studies.

### Predictive accuracy of DNA Embeddings

We trained Convolutional Neural Network (CNN) to optimize vector representations of the DNA barcodes in the Euclidean space. The model yielded impressive 99.44% accuracy on the holdout validation set that was created by reserving 20% of the training set. In the deployment of Neural Network models, it is also important to test the generalizability of the model to completely unseen classes/ species. Towards this end, we trained a K-Nearest-Neighbor (KNN) classifier (*K* = 1) on randomly sampled 80% of the DNA embeddings of unseen classes (243 species) obtained from our CNN model and tested on the remaining 20%. The simple KNN classifier rendered 99.19% accuracy proving the robustness of the CNN model to learning representation for undescribed species.

### Open-set Bayesian Classification with Unknown/Undescribed Species (OSBC)

No class information can be defined for undescribed species as these species are completely unrepresented in the training data. The only data available for training are images and DNA barcodes from described insect species (seen classes). The machine learning task at test time involves identifying insect classes originating from described species at the species level and those from undescribed species at the genus level. The Bayesian model is first trained and tested with CNN barcode embeddings (OSBC-DNA) and then with ResNet101 (36) image embeddings (OSBC-IMG). Finally, the CNN barcode and ResNet image embeddings were investigated jointly to determine if image information can improve the accuracy of the DNA Barcode classifier in inductive as well as transductive settings. As a standard approach to fusing DNA and image information in the inductive setting, the DNA and image embeddings were concatenated into a single feature vector (OSBC-DIC). Another approach in the inductive setting we tried was the summation of normalized likelihood vectors generated by two Bayesian classifiers of CNN and ResNet embeddings (OSBC-DIL). Finally, we developed a transductive approach that optimizes a linear mapping from image space to DNA sequence space by solving a ridge regression problem using ResNet and CNN embeddings of all available cases in test and train sets without using any class labels (OSBC-DIT). Additionally, we also developed a simple baseline using DNA sequences from the bioinformatics tool present in Matlab. Technical details of the Bayesian classifier, baseline method, and transductive approach are included in the Methods Section.

Table 1 reports the results from open-set insect classification. Once the number of classes reaches thousands, image classifiers alone cannot offer high performance emphasizing the need for high-quality 3D images. DNA data, on the other hand, proves to be very informative for species classification. The bioinformatics baseline method using DNA alone was excellent at accurately classifying seen species (species that are present in the database) while achieving an accuracy of 72% on unseen species, a significant reduction in comparison to using OSBC-DIT. Although OSBC-DNA yields a better unseen class accuracy, the performance on seen classes slightly drops.

**Table 1.** Open set classification results.

| Methods | US | S | H |
| --- | --- | --- | --- |
| OSBC-IMG | 35.88 | 39.11 | 37.42 |
| BioInformatics (DNA) baseline | 71.85 | **98.65** | 83.16 |
| OSBC-DNA | 73.39 | 96.15 | 83.24 |
| OSBC-DIC | 77.26 | 97.26 | 86.25 |
| OSBC-DIL | **81.95** | <u>98.21</u> | **89.35** |
| OSBC-DIT ($Tr + Ts_s + Ts_{us}$) | <u>81.39</u> | 96.66 | <u>88.37</u> |
| OSBC-DIT ($Tr + 50\%Ts_{us}$) | **79.94** | 96.66 | **87.53** |
| OSBC-DIT ($Tr + 25\%Ts_{us}$) | 77.48 | 96.63 | 86.01 |

US and S represent unseen and seen class accuracies and H represents the harmonic mean of these two scores. For both seen and unseen classes, each class accuracy is calculated then the average of these class accuracies is reported. Note that, these results are on genus level for unseen classes. More precisely, during class accuracy calculation different unseen classes belonging to the same genus are treated as the same class. Best results are displayed in bold and the second-best results are underlined. $Tr$ , $Ts_s$ and $Ts_{us}$ represents train, test seen and unseen data, respectively.

In all three scenarios, combining image and DNA data clearly helps the Bayesian classifier harvesting a decent performance boost, in particular for unseen classes. Transductive and heuristic likelihood methods perform the best with above 88% harmonic mean and 81% unseen class accuracy. That being said, both inductive methods, OSBC-DIC and OSBC-DIL, have an inherent flaw: they always require test samples to have an accompanying image along with a DNA barcode. For the transductive method (OSBC-DIT), however, only a fraction of test data with image and DNA pair, without using any labels, is enough to learn a robust mapping and still deliver a remarkable performance increase. The main information flow in learning the mapping in the transductive setting is coming from unseen classes. The last two rows of Table 1 display model performance while utilizing various fractions of image and DNA test data pairs from unseen classes for learning image to DNA embedding. Note that, we did not tune the model for these configurations and employed the validation parameters used to produce OSBC-DIT results. Using only 25% of image-DNA pairs from unseen classes to learn the Ridge regression already improves the harmonic mean to 86%. This finding clearly displays how the abundance of unlabeled image and DNA pairs can be leveraged by the transductive method to significantly boost the DNA alone classifier performance.

The transductive model (OSBC-DIT) yields 96.66% overall accuracy of seen class classification with 4,827/4,965 correct classifications (See Table 2). For unseen classes, the accuracy declines, unsurprisingly, but is remarkably good for 3 of the 4 orders with >80% accuracy of assigning the unknown “species” to the correct genus (with an overall accuracy of 81.01%). The order Diptera is where a large portion of unseen classes was misclassified to the incorrect genus (Table 2). When examining the different family groups and their classification accuracy (Table 2), the Culicidae (the mosquitos), Syrphidae (the hover flies), and Tipulidae (the crane flies) had the greatest amount of misclassifications, however, the number of possible species in the group is not accounting for the misclassifications, as species in Chironomidae were classified with 100% accuracy. With the Culicidae, 45/58 of the misclassifications were Aedes vexans records that classified to the Culex genus. When taking a random record and using the DNA sequence to BLASTn (37) in Genbank as a semi-independent test of the data, there are BOLD records that populate the hit list that are Culicinae sp., and therefore, these records may be obstructing the classification due to the overlap in sequences. For the Syrphidae, 18 Platycheirus *neoperpallidus* records were misclassified to Platycheirus *clypeatus*. When random P. *neoperpallidus* records were aligned to other Platycheirus species, it was noted that there was a great deal of similarity with P. *quadratus*, a species not present in the training set, again, demonstrating the need for a more representative training dataset to ensure accuracy within certain groups. There was only one instance in which every single individual was misclassified, wherein 14 records of the Tipulidae, all belonging to a single species Tipula *coloradensis*, were completely misclassified. The majority of the misclassifications were to the same subfamily (Tipulinae), but misclassified to the Nephrotoma genus, and four of the 14 misclassified to Syrphidae. What is remarkable with this dataset is that the training data contained three species of Tipula (T. *caliginosa*, T. *salicetorum*, T. *shirakii*). Sequence similarities were calculated between the three in the training set and T. *coloradensis*, and what is apparent very quickly is that T. *salicetorum* and T. *caliginosa* are closely related (interspecific sequence differences 97%), whereas the sequence similarity of T. *coloradensis* with either T. *salicetorum* or T. *caliginosa* is 88%. Further, T. *shirakii* is perhaps the most different, with 85% sequence similarities from the remainder of the Tipula species included in this analysis (data not shown). What this is indicative of is quite the vast amount of sequence variation that may exist in this genus.

**Table 2.** Seen and unseen class accuracy by insect family for five or more species per family.

| Order | Family | Seen Classes |  |  | Unseen Classes |  |
| --- | --- | --- | --- | --- | --- | --- |
|  |  | # training | # test samples | Accuracy | # test samples | Accuracy |
| Coleoptera | Brentidae | 94 | 18 | 100.00% |  |  |
|  | Cantharidae | 226 | 43 | 93.02% | 77 | 94.81% |
|  | Carabidae | 1660 | 346 | 95.66% | 128 | 95.31% |
|  | Cerambycidae | 210 | 43 | 100.00% |  |  |
|  | Chrysomelidae | 564 | 114 | 99.12% | 37 | 89.19% |
|  | Coccinellidae | 226 | 46 | 100.00% |  |  |
|  | Curculionidae | 348 | 68 | 94.12% | 55 | 96.36% |
|  | Dytiscidae | 146 | 30 | 100.00% | 18 | 88.89% |
|  | Elateridae | 242 | 47 | 100.00% | 12 | 100.00% |
|  | Scarabaeidae | 106 | 23 | 91.30% |  |  |
|  | Staphylinidae | 714 | 150 | 92.67% | 47 | 100.00% |
|  | Tenebrionidae | 186 | 24 | 100.00% |  |  |
| **Summary (C)** | **37** | **5,680** | **1,143** | **95.80%** | **751** | **85.22%** |
| Diptera | Calliphoridae | 190 | 35 | 100.00% | 13 | 92.31% |
|  | Chironomidae | 464 | 96 | 97.92% | 24 | 100.00% |
|  | Culicidae | 496 | 107 | 89.72% | 58 | 22.41% |
|  | Drosophilidae | 392 | 85 | 84.71% | 80 | 81.25% |
|  | Muscidae | 104 | 22 | 90.91% |  |  |
|  | Sciaridae | 150 | 33 | 100.00% |  |  |
|  | Syrphidae | 342 | 71 | 97.18% | 45 | 60.00% |
|  | Tipulidae | 122 | 26 | 96.15% | 14 | 0.00% |
| **Summary (D)** | **20** | **2,744** | **570** | **93.68%** | **273** | **61.17%** |
| Hymenoptera | Andrenidae | 192 | 39 | 100.00% | 53 | 79.25% |
|  | Colletidae | 190 | 32 | 100.00% | 56 | 100.00% |
|  | Crabronidae | 312 | 66 | 100.00% | 60 | 96.67% |
|  | Eulophidae | 226 | 47 | 100.00% | 183 | 100.00% |
|  | Halictidae | 344 | 70 | 98.57% | 113 | 80.53% |
|  | Ichneumonidae | 306 | 67 | 100.00% | 12 | 100.00% |
|  | Megachilidae | 296 | 55 | 100.00% | 28 | 53.57% |
|  | Tenthredinidae | 864 | 169 | 91.72% | 261 | 66.28% |
|  | Vespididae | 106 | 22 | 100.00% | 22 | 77.27% |
| **Summary (H)** | **19** | **3,282** | **660** | **97.27%** | **872** | **82.22%** |
| Lepidoptera | Coleophoridae | 994 | 206 | 99.51% | 170 | 82.35% |
|  | Crambidae | 1054 | 176 | 99.43% | 482 | 87.14% |
|  | Depressariidae | 1836 | 269 | 100.00% | 380 | 67.63% |
|  | Erebidae | 4288 | 464 | 97.20% | 694 | 74.78% |
|  | Gelechiidae | 268 | 59 | 96.61% | 41 | 82.93% |
|  | Geometridae | 1170 | 230 | 96.96% | 328 | 89.63% |
|  | Hesperiidae | 2294 | 14 | 85.71% | 566 | 47.00% |
|  | Noctuidae | 3246 | 570 | 98.95% | 525 | 82.10% |
|  | Notodontidae | 4068 | 257 | 100.00% | 959 | 94.89% |
|  | Nymphalidae | 554 | 37 | 100.00% | 166 | 84.94% |
|  | Saturniidae | 890 | 31 | 100.00% | 111 | 99.10% |
|  | Tortricidae | 968 | 170 | 100.00% | 144 | 96.53% |
| **Summary (L)** | **18** | **22,564** | **2,592** | **98.61%** | **6,567** | **81.01%** |

Note that ‘Summary’ row reports the summary results from all families belonging to that order including families having less than five species in our dataset.

## Discussion

Deep learning methods are getting more and more integrated into various fields and disciplines in the sciences. Here we present a novel method for potential classification of new insect species, with an eye on the future of identification through image analysis and character extraction for the entomology field explicitly, although this can be applied to any biological organism. This is the first attempt where an open set classification is done using the integration of DNA and image analysis on a comparatively larger number of classes (in this case, 1,040 species in four large orders). The use of image analysis alone, or DNA analysis alone, has had varying levels of success. DNA is generally viewed as strong support for new species if the sequence variation falls outside the normal bounds of intraspecific variation, however, those normal bounds can vary significantly. In some cases, the DNA barcode has been integral to differentiate between species that are morphologically indistinguishable, confirmed through additional nuclear DNA sequencing (38). Image analysis alone has provided some gains in order to monitor (in real-time) insect species but suffers when background extraction is necessary. Furthermore, these methods are closed-set since the application is related to monitoring for existing species (for example, when pest management strategies are necessary) (39, 40). When using deep learning methods with images to identify seen classes of insects, accuracy gains can reach 90 percent or higher (29-igher (29–31, 41), in some cases, approaching or surpassing taxonomic specialist accuracy’s (42). However, all these methods are tested either on coarse-grained datasets or with a very limited number of classes, generally less than 15 species. Furthermore, the lingering issue of identifying unseen classes and the inherent data imbalance continue to plague the ability of more efficient means of identifying new species, especially within the Insecta class, where the majority of the species continue to be unidentified and presents the most important advancement to the field of entomology, but more broadly, to better understanding ecosystems and their processes, of which insects likely play a major role (43). The model trained on DNA embeddings (OSBC-DNA) achieved a compelling 96.15% accuracy on seen classes where 670 out of 770 test classes are perfectly classified to their true species. The model performance dropped to 73.39% in a more challenging task of identifying unseen species and assigning them to their true genera. OSBC-DNA completely misclassified all samples of 24 unseen species (less than 10% of all unseen classes), yet it is worth noting that 6 of these classes were perfectly assigned to their true classes as the second-best option. Leveraging the auxiliary image data, our transductive approach (OSBC-DIT) significantly boosts the unseen class performance to 81.39% (and 11% increase) with a modest increase on the seen class accuracy over DNA alone (OSBC-DNA). OSBC-DIT classified 677 out of 770 seen classes with 100% accuracy. The model also partially recovered 14 of 24 completely missed unseen species under OSBC-DNA model (see Fig 2), where 9 out of 14 classes were recovered by more than 80%.

**Fig. 2.**
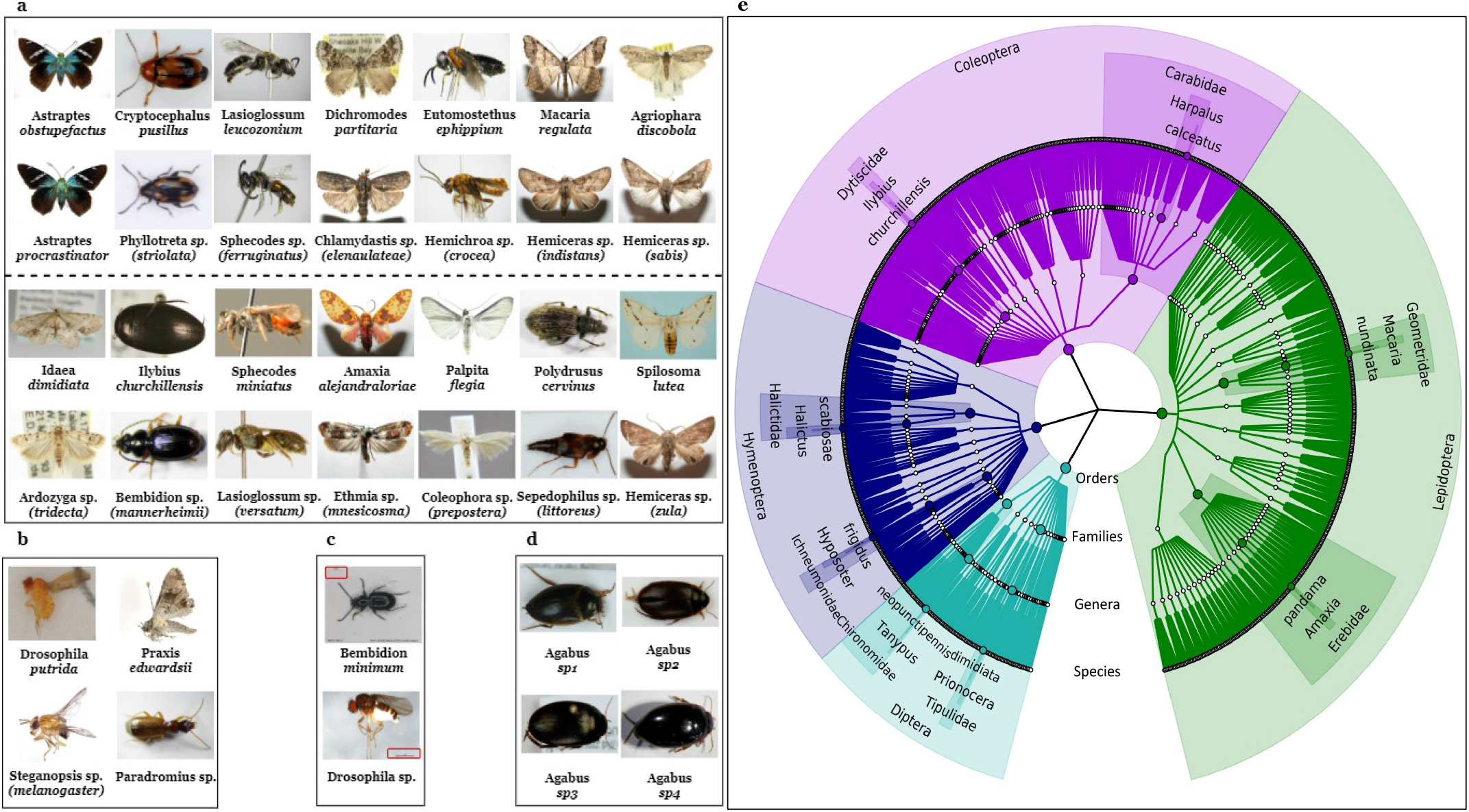
Discussion cases and phylogenetic tree. **a**. Unseen classes (14) that were completely missed by OSBC-DNA but discovered by OSBC-DIT (some species partially and some fully). The first and third rows display the images that are covered by OSBC-DIT and the second and fourth rows show samples for the corresponding classes that they are misclassified to under OSBC-DNA. Names containing “sp.” means that this is a Genus class and the image is from the species of name in parenthesis belonging to that genus. **b**. Misclassified cases due to image manipulation. **c**. Misclassified case due to background noise. **d**. Morphological resemblance between species belonging to the same genus. **e**. Phylogenetic tree of the 4 orders from the dataset. Randomly chosen 2 species from each order with their full taxonomic order are illustrated.

### Striking morphological similarity between species belonging to the same genus

As it is observed in Figure 2a, variation in some insects is nearly invisible to the human eye, especially if lacking specialized expertise, yet the models were able to extract these subtle differences from images and aid DNA embeddings to correctly classify these difficult cases. To illustrate, we present a simple challenge in Fig 2d where one sample from 4 different species belonging to Agabus genus is displayed. The task is to correctly match the images with the following species names: A. *sturmii*, A. *bipustulatus*, A. *uliginosus*, and A. *infuscatus*. The true order can be found in this footnote^*^. Out of four species, A. infuscatus is reserved as an unseen class. DNA classifier correctly classified all test samples from 3 seen classes, however, it made a few mistakes while assigning the samples of unseen class into its true genus, Agabus. OSBC-TID model, on the other hand, correctly classified with 100% accuracy all seen and unseen class test samples.

This observation also reveals that 658bp of DNA sequence (cytochrome oxidase subunit I) can miss some subtle features, yet image representation can highlight these features as spotted in the Lasioglossum and Sphecodes cases (column 3 of Fig 2a). Both genera share very similar DNA sequences and are members of the same tribe (Halictini), which makes it quite difficult to differentiate using merely DNA barcodes in a challenging open-set classification setup. In the transductive approach (OSDB-DIT), these elusive morphological features are successfully transferred from image space to DNA space and fill the gap in the utility of DNA barcodes.

### Effect of image quality and background noise on model performance

High-quality images are an integral part of any successful machine learning approach and heavily impact the model performance in computer vision tasks. It is well documented in the literature that due to cross-entropy loss they have been trained with, many state-of-the-art pre-trained CNN models are sensitive to the presence of subtle noises such as Gaussian, background noises, or blurriness in the image (44–46). The following interesting cases observed in our experiments also verified these phenomena where a few isolated instances were misclassified to unrelated species under OSBC-DIT classifier.

Cases in Figure 2b illustrate the vulnerability of the CNN models towards image manipulations. The first case is a test sample from the seen class Drosophila *putrida*, which is correctly classified by the DNA classifier, yet OSBC-TID misclassified the sample to Steganopsis genus. Except for this case, all the cases from seen classes where the DNA model correctly classified but DNA+IMAGE model failed were either misclassified to the true genus or to another species from the corresponding genus. In the light of this statistic, this particular case stands out as the OSBC-TID model misclassified this test sample (and only misclassified test case from D. *putrida*) to another genus. Inspecting the image features reveals that this figure is the only one being exposed to image manipulation and has been trimmed by Adobe Photoshop CS (this information can be accessed from image properties). In the same fashion, the test case from P. *ewardsii* species in Figure 2b is the only test sample that was misclassified by OSBC-TID, and also the only sample exposed to a modification from a software called CombineZP (47) (this information can be accessed from image properties). These subtle alterations are most of the time indistinguishable to human eyes yet can drastically alter the CNN model embeddings. Recent research suggests more robust image embeddings less sensitive to subtle alterations can be obtained using backbone architectures trained by self-supervised learning (44, 48). Background information can sometimes dominate the relevant image features. The aberrant misclassification of the test sample from Bembidion minimum to a Drosophila genus (from different order) is an example of this phenomenon (See Figure 2c). Many images from Drosophila genus have “1mm” text attached next to the species image to illustrate the scale, and that particular test sample (the only misclassified sample from B. *minimum*) has the same “1mm” text in the background.

## Conclusion

All living beings, including plants, have a complex and intertwined relationship contributing to the delicate balance our planet has been maintaining. There have been drastic changes observed in the last few decades, disturbing this balance. These alterations reflect their consequences first on biodiversity, thus it has vital importance to measure and monitor these effects. In this study, we developed a novel framework to facilitate the discovery and identification of insect species, a very large swatch of unknown biodiversity, at scale. The proposed model is the first in the literature to tackle this problem by leveraging the image and DNA data together tested on more than a thousand species. Unlike all the previous work, our model does not simply cast aside the new insect species by treating them as an outlier but classifies them to the lowest level of abstraction in taxonomic order, the genus level. Our transductive Bayesian classifier delivered 81% accuracy on identifying the correct genus of new species that have no image or DNA samples present in our training data, meanwhile classified known species with more than 96% accuracy. Considering the transductive approach was built on regularized linear mapping, it appears there is a great potential to achieve better performance utilizing nonlinear and more sophisticated approaches like Generative Adversarial Networks (49) or Variational Autoencoders (50) to learn this mapping. Integrating GAN/ VAE would also allow training an end-to-end model by self-supervised learning that can potentially mitigate the shortcomings of supervised pre-trained models.

In this proof of concept, the focus of this paper was on new species discovery, wherein the subclasses were species, and the superclasses were genera. The Bayesian model can easily be extended to be trained on where genera/species are considered the subclasses and higher taxonomic levels are considered superclasses (e.g., family). Such a classifier will readily deal with missing/unobserved genera. However, a more extensive dataset covering more genera and families would be necessary.

## Materials and Methods

In this section, we first introduce the dataset and how the split is performed for machine learning training. Next, Convolutional Neural Network (CNN) model for deriving DNA embeddings is presented. Finally, we lay out the open-set Bayesian classifier details along with the bioinformatics baseline classifier.

### Barcode of Life Data System

Our study uses insect data from the Barcode of Life Data System (BOLD) (12, 13). As other databases exist of genetic data (for example, (51)), they require some identification prior to depositing into the database. BOLD differs slightly in that as it allows for unidentified organisms to be uploaded into the database, and their algorithms, based on DNA sequence only, will place the unknown into a barcode index number (BIN). This allows for the quantification of the unknown and undescribed, however, no identifications are made. This data repository does not contain samples of truly undescribed species. The BOLD database using a specific searching algorithm that translates the DNA sequence to its protein sequence and searches its database. BOLD will make a species identification if the queried sequence contains less than 1% divergence to a reference specimen located in the database. If the sequence divergence is less than 3% (but greater than 1%), the database will make a match to a genus.

All insect image and DNA sequence pairs in our dataset were downloaded from the Barcode of Life Data System. Most insects in the database had approximately 658bp of the DNA barcode (cytochrome oxidase subunit I), as well as an image and additional information such as country of origin, life-stage, order, family, sub- family, and genus/species names.

BOLD is an open-access database in which users can upload DNA sequences and other identifying information regarding any animal on Earth. Because the majority of the uploads are not identified species, they are classified into BINs (13). For example, as of 8/18/2021, the Insecta database had a total of 5,883,100 records with sequences, and about half had species names (2,561,685), meaning the remainder could not be identified to species. The data are important for assessing biodiversity, distributions of species, as well as collating other descriptive metadata and images. The limitations of this database are that it is important for the discovery of new species but does not allow for the identification of such, and simply places the outliers in an interim position, not allowing for any forward movement.

### Data Collection

Data were collected based on a subset of insects that originate from four major Insecta orders: Diptera (true flies), Coleoptera (beetles), Lepidoptera (butterflies and moths), and Hymenoptera (sawflies, wasps, bees, and ants). While the dataset was generally clean, manual effort was devoted to further curate the dataset. Only non-teneral adults with images and matching DNA barcodes were included with each species and manually inspected so that images with low quality, duplicates, images with just insect parts, or missing images (e.g. just a label is present) were deleted. Only classes that had a minimum of 10 images within a single BIN were included in the final dataset. Consequently, the final dataset consisted of 1,040 insect species and a total of 32,848 insect instances (records). In the finalized dataset, we obtained 108 species of Diptera from 63 genera, 329 species of Coleoptera from 164 genera, 189 species of Hymenoptera from 59 genera, and 414 species of Lepidoptera from 82 genera (See Figure 2e)

A pre-trained ResNet101 model (36), 101-layered Convolutional Neural Network, was used to embed images into Euclidean vector space and represented them by information-rich 2048 dimensional real-valued feature vectors. We utilized the ResNet101 model parameters that were optimized on ImageNet 1000 classes, hence pre-trained, and we did not fine-tune the model on our dataset. Images are first resized to 256×256, then center-cropped into the ResNet model image dimension: 224×224. No other pre-processing was applied to the images.

### Split details

The BOLD database does not contain truly undescribed species. To artificially create undescribed test classes, genera were chosen that have a minimum of three species, and 33% of those species were randomly chosen and set aside as undescribed species. These pseudo-undescribed species are referred to as unseen classes and described species as seen classes. For example, the genus Coelioxys has three species, and one of them (in our case C. *conoidea*) was randomly chosen as an undescribed species, leaving two as seen classes. This split left 243 unseen classes and 797 seen classes, where the training set did not include any images or DNA from these 243 classes. In order to create a validation set for unseen classes, in the same fashion, 33% of species were randomly chosen of genera that have at least three members from the 797 training classes. The remainder of the data were split by a 70/30 ratio in a stratified fashion to obtain samples for training and test seen classes. Some of the insect classes have multiple images, each capturing a different view of the insect (for example, ventral and dorsal views), all insect classes with multiple images were restricted to the training set, leaving 27 of the seen classes with no available samples for testing. Test samples from seen and unseen classes summed up to 4,965 and 8,463 instances.

### CNN Embeddings for DNA Barcodes

A Convolutional Neural Network (CNN) (52, 53) was trained to optimize vector representations of the DNA barcodes in the Euclidean space. Barcodes are first converted into 658×5 2D one-hot encoding arrays, where 658 is the length of the barcode sequence (median nucleotide length of the DNA data). A total of five tokens were used, one for each of the Adenine, Guanine, Cytosine, Thymine bases, and others. All ambiguous and missing symbols are included in the others token. To train the CNN model, a balanced set out of the training data, which was discussed in the previous paragraph, is created, where class sizes are capped at 50 samples. The training set is finalized with 14,389 barcodes from 797 classes. Note that no barcodes nor images from 243 unseen classes or test data are employed during model training. The training set is further split into two sets as the train (80%) and validation (20%) by random sampling. We used 3 blocks of convolutional layers each, followed by batch normalization and 2D max-pooling. The output of the third convolutional layer is flattened and batch normalized before feeding the data into a fully-connected layer with 500 units. The CNN architecture is completed by a softmax layer. For the embeddings, we used the output of fully-connected layer. The details of the model architecture are depicted in Supplementary Fig 4. We trained the model for 5 epochs with a batch size of 32 and used Adam optimizer (54) (learning rate = 0.0005 and drop factor= 0.5, *β*_1_ = 0.9, *β*_2_ = 0.999). The model is developed in Python with Tensorflow-Keras API.

### Open-set Insect Classification

In our approach, we assume that there are species that are completely unknown (for example, a newly discovered species), and we introduce a framework that can identify insects at the lowest taxonomic level possible by jointly leveraging image and taxonomic information. More specifically, if an insect to be classified is a previously described species, the test sample would be classified as one of the species present in the training set. On the other hand, if the insect is undescribed and therefore not present in the training data, the taxonomic level identification would be to genus, providing clues that the insect is not a species in the current database. Thus, for undescribed insect species, the genus would be predicted, therefore indicating the database/training does not contain the species and it is likely an unknown species. This open-world classification approach not only significantly reduces the uncertainty surrounding traditional closed-set supervised algorithms (closed-set algorithms assume all possible classes/ species are present in the training data and therefore would misclassify all new/ undescribed insects into one of the known species), but also addresses problems with existing open-set frameworks where any undescribed species are designated as an outlier, thus no additional taxonomic level is being identified. We explore the predictive accuracy of the open-world classifier first by DNA barcodes alone, then by images alone, and finally by combining DNA barcodes and images in various forms.

### Bayesian Model

Insect species have a predefined taxonomic hierarchy; species < genus < subfamily < family < order etc., although rich variety between these hierarchies carries valuable information, it is often overlooked when designing ML algorithms. A hierarchical Bayesian model was recently introduced in computer vision for zero-shot classification of object classes (55). This model establishes a Bayesian hierarchy among object classes using visual attributes as auxiliary information. To identify both described and undescribed species a similar model is developed by replacing visual attributes with a predefined class hierarchy explicit in the taxonomical classification of biological organisms. More specifically, our proposed method assumes that there are local priors that define the class hierarchy in the feature space (image or DNA) and uses predefined taxonomical classification to build the Bayesian hierarchy around these local priors. Supplementary Fig 2 illustrates the intuition behind this idea: species sharing similar haplotypes cluster in the phenotypic space as well. Our model uses two types of Bayesian priors: global and local. As the name suggests, global priors are shared across all species, whereas local priors are only shared among species belonging to the same genus. Unlike standard Bayesian models where the posterior predictive distribution (PPD) establishes a compromise between prior and likelihood, our approach utilizes posterior predictive distributions to blend local and global priors with data likelihood. Inference for a new insect sample (image or DNA) is performed by evaluating these posterior predictive distributions and assigning the insect to one of the described species that maximizes the posterior predictive likelihood or identifying it as a new species belonging to the surrogate genus class maximizing the posterior predictive likelihood.

### Generative model

The Supplementary Figure 3 depicts the graphical model of the proposed approach with the model design given below:

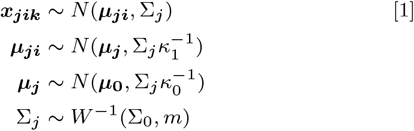

where *j, i, k* represent indices for local priors, classes, and data instances, respectively. We assume that the instance ***x***_***jik***_ comes from a Gaussian distribution with mean ***µ***_***ji***_ and covariance matrix Σ_*j*_. They are generated independently conditioned not only on the global prior but also on their corresponding local priors.

Each local prior is characterized by the parameters ***µ***_***j***_ and Σ_*j*_. ***µ***_**0**_ is the mean of the Gaussian prior defined over the mean vectors of local priors, *κ*_0_ is a scaling constant that adjusts the dispersion of the centers of local priors around ***µ***_**0**_. A smaller value for *κ*_0_ suggests that class centers are expected to be farther apart from each other whereas a larger value suggests they are expected to be closer to each other. On the other hand, Σ_0_ and *m* dictate the expected shape of the class distributions, as under the inverse Wishart distribution assumption the expected covariance is 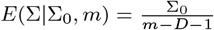, where *D* is the dimension of feature space. The minimum feasible value of *m* is equal to *D* + 2, and the larger the *m* is the less individual covariance matrices will deviate from the expected shape.

The hyperparameter *κ*_1_ is a scaling constant that adjusts the dispersion of the class means around the centers of their corresponding local priors. A larger *κ*_1_ leads to smaller variations in class means compared to the mean of their corresponding local prior, suggesting a fine-grained relationship among classes sharing the same local prior. Conversely, a smaller *κ*_1_ dictates coarse-grained relationships among classes sharing the same local prior. In this model, classes sharing the the same local prior also retain the same covariance matrix Σ_*j*_ to preserve conjugacy of the model. Test samples are classified by evaluating posterior predictive distributions (PPD) of seen and unseen classes.

### PPD derivation

PPD incorporates three sources of information: the data likelihood that arises from the current class, the local prior that results from other classes sharing the same genus as the current class, and global prior defined in terms of hyperparameters. The derivation in six steps are outlined in Supplementary Figure 3a and Algorithm 1 describes a pseudo code on deriving PPD for both seen and unseen classes. Class sufficient statistics are summarized by 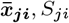 and *n*_*ji*_ which represent sample mean, scatter matrix and size of class *i* of local prior *j*, respectively.

PPDs for seen classes include the global prior and data likelihood (See Fig 3) and are derived in the form of a Student-t distribution as below,

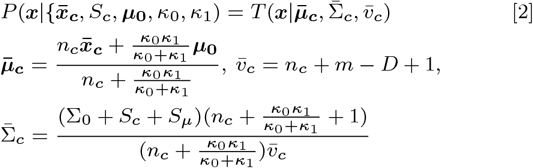

where, 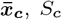 and *n*_*c*_ are sample mean, scatter matrix and size of current seen class *c. S*_*µ*_ is defined in Equation (34) from Supplementary material. The index *c* in Equation (2) represents the current seen class, whose PPD is being derived.

**Fig. 3.**
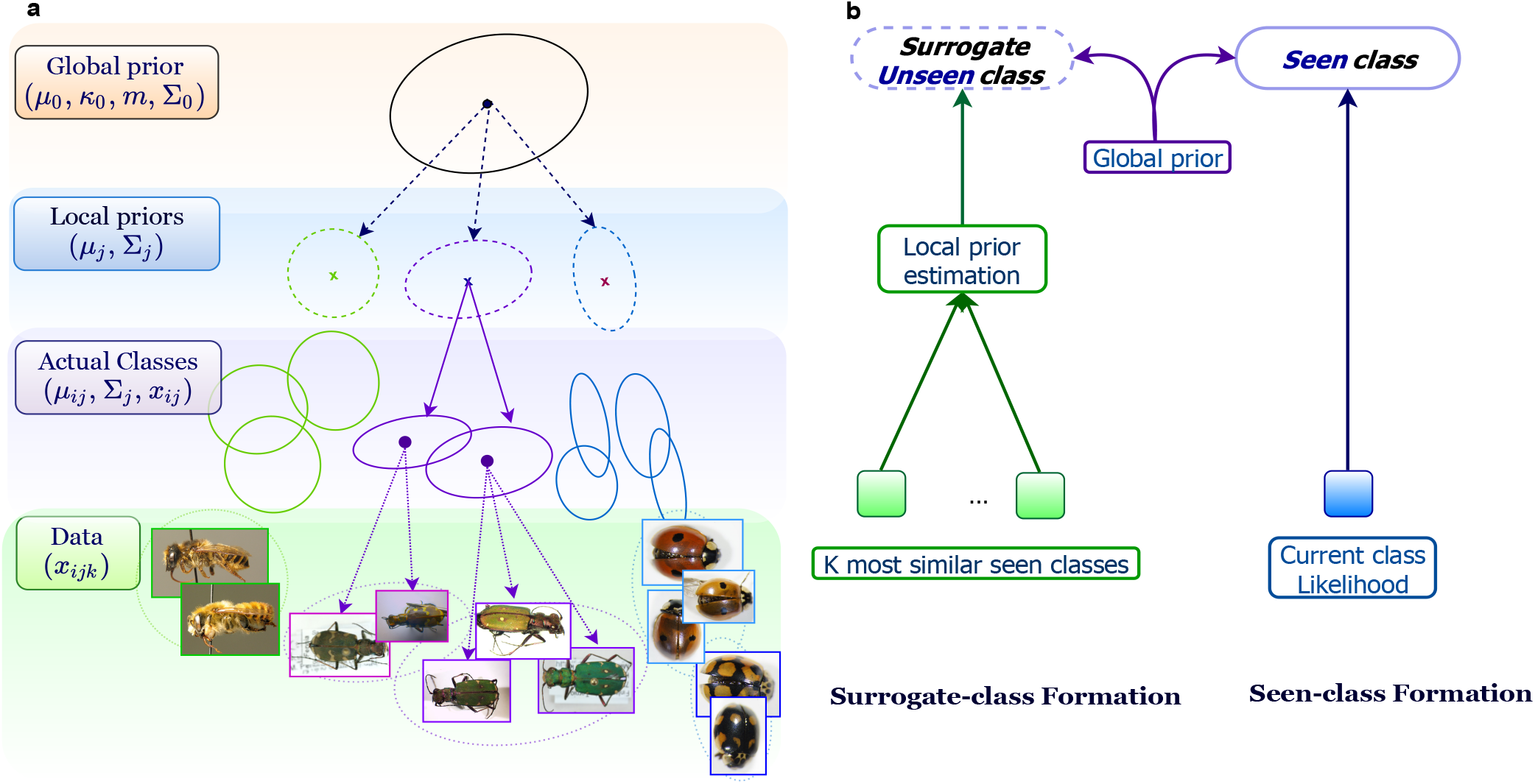
**a**. Generative model. Hyperparameters are defined in the Methods section. **b**. Class distribution formation for seen and surrogate genus classes.

#### Algorithm 1 Modeling seen and surrogate genus classes in BZSL

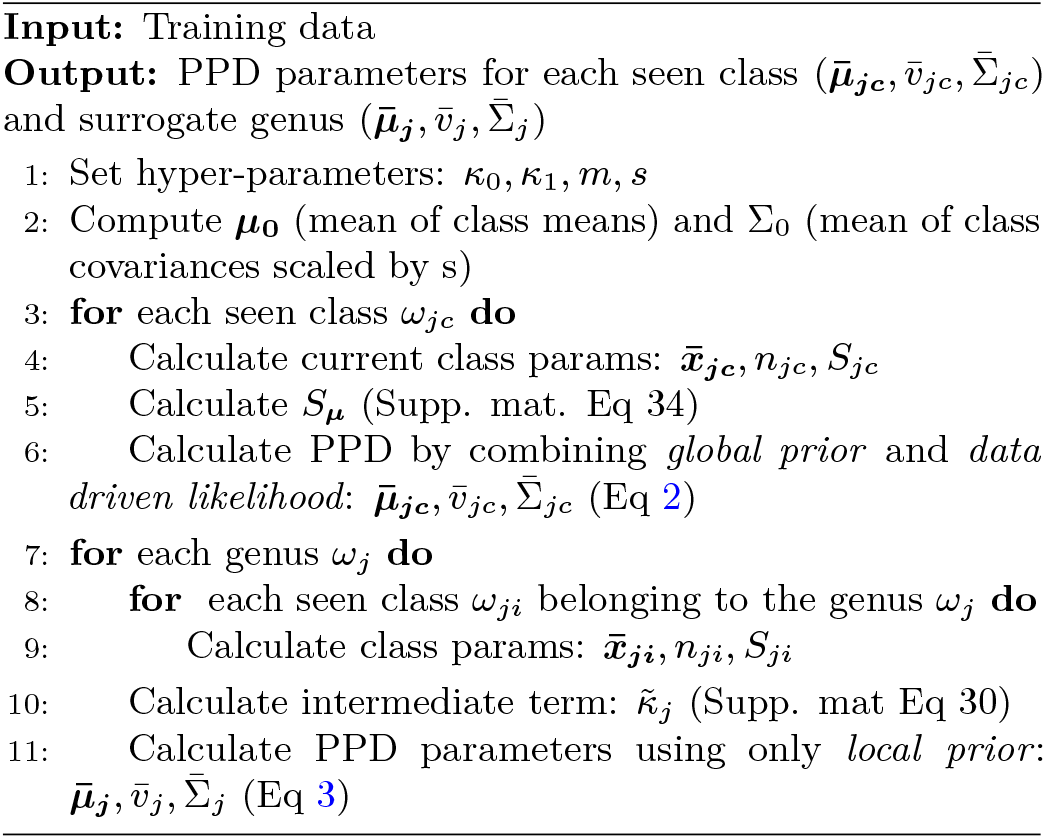

### Surrogate class formation

In our model, groupings among classes are based on local priors. Hence, once estimated from seen classes, local priors can be used to define surrogate classes for unseen classes during inference. We form a surrogate-class for each genus in our dataset by forming a local prior combining all seen classes from that genus (See Fig 3b). During the inference, test samples are classified based on class-conditional likelihoods evaluated for both seen and genus-level surrogate classes.

PPDs for unseen classes also follow a Student-t distribution, thanks to conjugacy, given below,

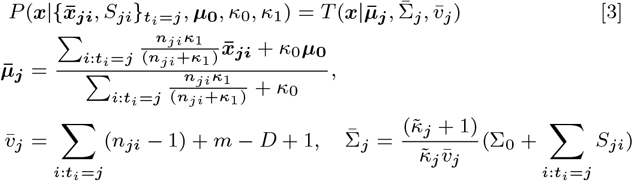

where, 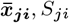 and *n*_*ji*_ represent sample mean, scatter matrix and size of class *i* associated with surrogate-class *j*, respectively and 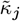 is defined as in Eq. (30) in the Supplementary material.

It is worth to clarify the distinction between seen and surrogate class PPDs in the case of genera where they have only one species in the training data. The seen class distribution and surrogate genus class will have similar formulas but with 2 important distinctions. First, mean of the seen class PPD will have more weight on class sample mean whereas mean of the surrogate class will lean towards *µ*_0_. Beside the common terms in location parameters, seen class PPDs have the term 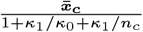 whereas surrogate class PPDs have *µ*_0_ in replace of 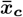. Second, unlike surrogate class PPDs, seen class PPDs have additional term, *S*_*µ*_, in their scale matrix.

### Transductive Approach

The transductive approach leverages the unlabeled test data as well during the training process. We aim to learn a linear mapping from Image feature space to DNA feature space using Ridge regression. Figure 1 panel (c) outlines the transductive approach. Following the notation in the figure, 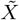 and *X* represents the image and DNA embeddings, respectively. 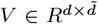 is the embedding from image space to DNA space we want to learn and *λ* is the regularization constant. Ridge regression with *Frobeneus* norm has a well-known closed for solution given as, 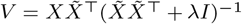. We leverage the learned mapping to augment auxiliary training data by embedding image features with labels into DNA feature space, mathematically 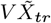 and combine this data with DNA embeddings. The whole process takes two lines of a code and computational cost is infinitesimal compared to the model training time, thus this step comes as free. Nonetheless, we achieve remarkable 10% percent performance boost on unseen class accuracy while preserving seen class accuracy.

### An Open-set Distance-based Bioinformatics Approach as a Baseline

For each described species, nucleotide sequences were aligned using training samples available for that species. Aligned sequences are then used to compute a consensus nucleotide sequence for each described species. Test samples were classified by evaluating Jukes-Cantor distance (56) between a test sequence and consensus sequences of described species. Test samples are assigned to the species with the minimum distance only if the minimum distance is smaller than a designated threshold. If the minimum distance is larger than this threshold then the test sample is treated as a sample of an undescribed species and assigned to the genus of the species with the minimum distance. Result of this approach is included in Table 1.

## Data and Code

The data and code can be accessed from [GitHub](https://github.com/sbadirli/Open-set-Insect-Discovery).

## ACKNOWLEDGMENTS

M.D. and S.B. were sponsored by the National Science Foundation (NSF) grant IIS-1252648 (CAREER). G.M. was sponsored by NSF grant ATD-2124313. The content is solely the responsibility of the authors and does not necessarily represent the official views of NSF.

## Footnotes

* A. *infuscatus*, A. *sturmii*, A. *bipustulatus*, and A. *uliginosus*

