## Supplementary material for "Classifying the Unknown: Identification of Insects by Deep Open-set Bayesian Learning"

Sarkhan Badirli
Department of Computer Science
Purdue University
West Lafayette, IN 47906

Christine J. Picard
Department of Biology
Indiana University - Purdue University
Indianapolis, IN, USA

George Mohler
Computer and Information Science Department
Indiana University - Purdue University
Indianapolis, IN, USA

Zeynep Akata
Computer Science Department
University of Tübingen
Baden-Württemberg, Germany

Murat Dundar
Computer and Information Science Department
Indiana University - Purdue University
Indianapolis, IN, USA


Although the main text is prepared to be self-contained, we provide further details for mathematical derivations, implementation, hyperparameter tuning, and limitations of the model.

### 1 Overview

#### 1.1 BOLD datasets

Although Insect data is complied from BOLD for this study, the database contains samples both image and DNA from many other animal groups as well as plants.

Figure 1 presents further details on INSECT dataset such as number of species per genera and number of samples per species. From Figure 1(a), one can observe that dataset can be considered balanced as around $90\%$ of species have samples between 10 and 50. Fine-grained nature of the dataset, on the other hand, can be seen from Figure 1(b). Out of 368 genera, more than half have at least 2 species in the dataset. In 65 genera, there are more than 4 species coming from the same genus, which makes the data challenging yet, at the same time, provides a chance to find similar seen classes for each unseen class and efficaciously model surrogate classes by the Bayesian classifier.

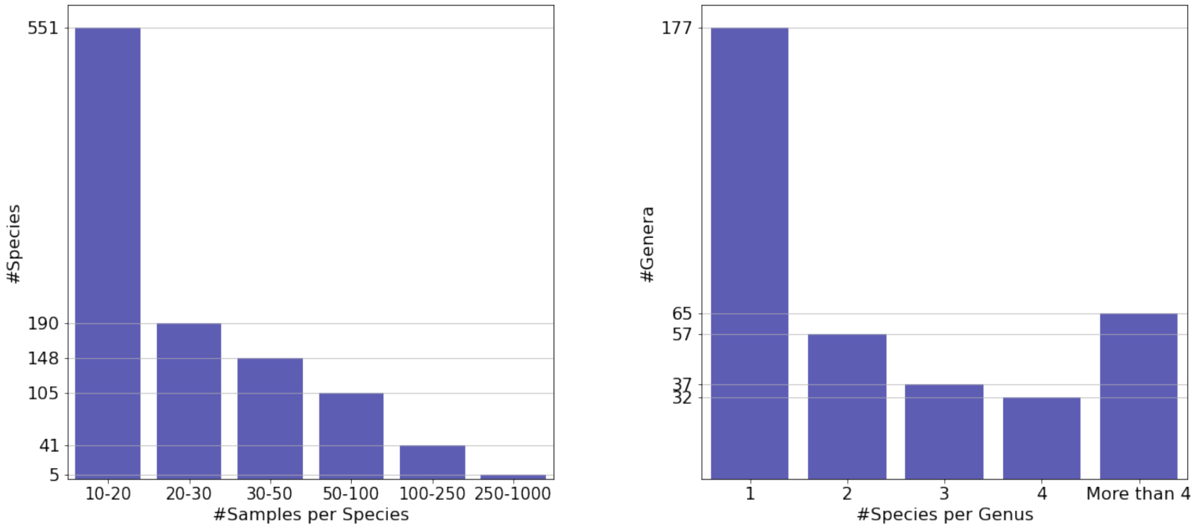


(a) Number of samples per species

(b) Number of species per genera

Figure 1: INSECT Data statistics.

#### 1.2 DNA embeddings and model intuition

Although traditional sequencing methods [2, 1] have shown that DNA barcodes extracted from COI gene are very powerful to uniquely identify living organisms in species level, it was to our surprise to see that DNA embeddings obtained from a simple CNN model yielded more than 99% accuracy on INSECT dataset with $1,040$ species. Figure 2 displays the T-SNE plot of DNA and image embeddings of species from 5 genera. On one hand, species clusters are well-separated. On the other hand, embeddings of species belonging to the same genus, are nicely clustered together. Note that, DNA embeddings of species belonging to the same genus, are very tightly clustered as such that only a single point appears for the whole cluster (Fig 2a). Image embeddings whereas are more loosely clustered (Fig 2b). Moreover, we can observe that 2 different genera from Hymenoptera clustered closely, in particular image space. These findings validate the model intuition that there is a deeper level of hierarchy among existing classes than a single global Bayesian prior can explain.

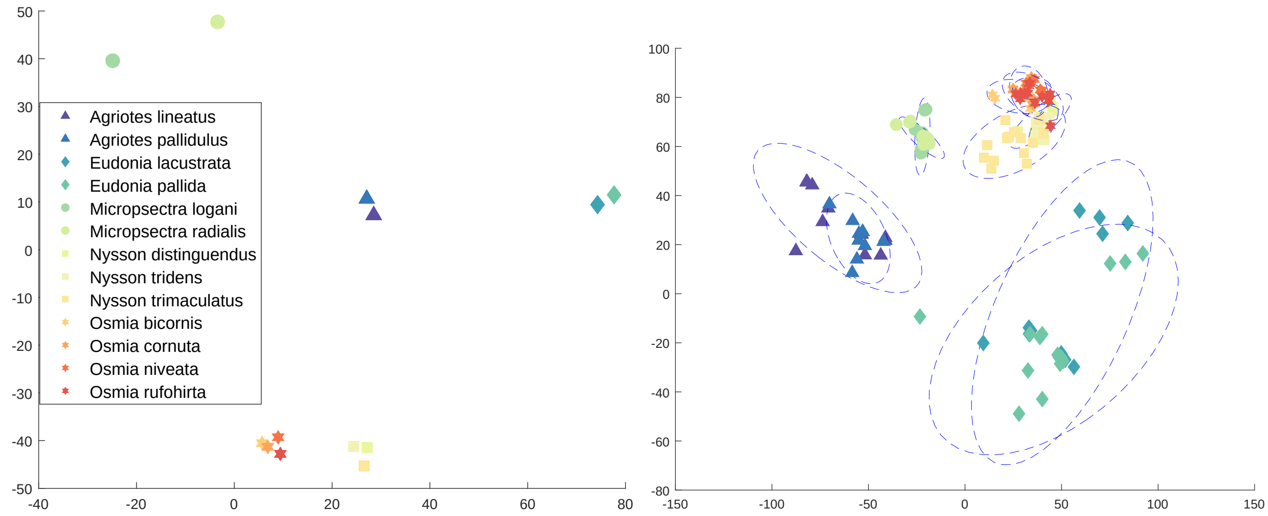


(a) Insect DNA embedding visualization in 2D

(b) Insect image embedding visualization in 2D

Figure 2: TSNE visualization of five genera, Agriotes (Coleoptera), Eudonia (Lepidoptera), Micropsectra (Diptera), Nysson, and Osmia (Hymenoptera) from the curated dataset. DNA embeddings are derived from our CNN model, and for image embeddings, we used pretrained ResNet101 model. Dashed contours are derived from class covariance matrices and placed at two standard deviations away from class means.

Furthermore, incorporating distinctive features of both feature spaces, as shown in the transductive approach from the main text, helps the model to achieve better performance.

### 2 Details of posterior predictive distribution (PPD) derivation

The proposed model (BZSL) assumes Gaussian data model and Normal-Inverse Wishart (NIW) prior on class parameters. Derivations for each class (seen and surrogate genus class) are treated independently as our model imposes independence on data generation conditioned on local priors and global hyperparameters. By preserving the conjugacy, the assumption of common covariance among classes sharing the same local prior further simplified the derivations. Six Steps of the derivation are outlined in Figure 3 and pseudo-code is presented in Algorithm 1. Please refer to the Table 1 for all variables and parameters used in the calculations. Class sufficient statistics, which are only available for seen classes, are defined by $\bar{\boldsymbol{x}}_{\boldsymbol{ji}}$, $S_{ji}$ and $n_{ji}$, which represent sample mean, scatter matrix and size of class $i$ associated with local prior $j$, respectively. The notations $\omega_{jc}$ and $\omega_j$ used in Algorithm 1 represents current seen and surrogate-genus classes, whose PPD are being derived.

Although we haven't utilized local priors for seen class, please note that the following derivations are based on a more general setup, where seen classes PPD can also incorporate local priors. At the end of the step 6, we will provide the exact PPD formulation that we used in our experiments in which local priors are omitted in Seen class PPD.

**Algorithm 1** Posterior Predictive Distributions in BZSL

**Input:** Training data
**Output:** PPD parameters for each seen class ($\bar{\boldsymbol{\mu}}_{\boldsymbol{jc}}, \bar{v}_{jc}, \bar{\Sigma}_{jc}$) and surrogate class ($\bar{\boldsymbol{\mu}}_{\boldsymbol{j}}, \bar{v}_j, \bar{\Sigma}_j$)

Set hyper-parameters: $\kappa_0, \kappa_1, m, s$
Compute $\boldsymbol{\mu_0}$ (mean of class means) and $\Sigma_0$ (mean of class covariances scaled by $s$)
**for** each seen class $\omega_{jc}$ **do**
    Calculate current class params: $\bar{\boldsymbol{x}}_{\boldsymbol{jc}}, n_{jc}, S_{jc}$
    Calculate $S_{\boldsymbol{\mu}}$ (Eq 34)
    Calculate PPD by combining *global prior* and *data driven likelihood*: $\bar{\boldsymbol{\mu}}_{\boldsymbol{jc}}, \bar{v}_{jc}, \bar{\Sigma}_{jc}$ (Eq 37)
**end for**
**for** each surrogate class $\omega_j$ **do**
    **for** each seen class $\omega_{ji}$ belonging to the genus $\omega_j$ **do**
        Calculate class params: $\bar{\boldsymbol{x}}_{\boldsymbol{ji}}, n_{ji}, S_{ji}$
    **end for**
    Calculate $\tilde{\kappa}_j$ (Eq 30)
    Calculate PPD parameters using *local* and *global priors*: $\bar{\boldsymbol{\mu}}_{\boldsymbol{j}}, \bar{v}_j, \bar{\Sigma}_j$ (Eq 38)
**end for**

| **Parameter** | **Description** |
|---|---|
| $D$ | image feature space dimension |
| $d$ | superscript that refers to the $d^{th}$ component of each parameter |
| $j$ | local prior index |
| $i$ | actual class index |
| $k$ | sample index |
| $c$ | index of current class |
| $t_i$ | local prior indicator for class $i$ |
| $t_k$ | class indicator for data point $k$ |
| $\boldsymbol{\mu_j}$ | mean of local prior $j$ |
| $\Sigma_j$ | covariance of local prior $j$ |
| $\boldsymbol{\mu_{ji}}$ | mean of class $i$ of local prior $j$ |
| $\bar{\boldsymbol{x}}_{\boldsymbol{ji}}$ | sample mean of class $i$ of local prior $j$ |
| $S_{ji}$ | scatter matrix of class $i$ of local prior $j$ |
| $n_{ji}$ | size of class $i$ of local prior $j$ |
| $\boldsymbol{x_{jik}}$ | data point $k$ from class $i$ of local prior $j$ |

Table 1: The notation used in the derivation of PPD.

#### Sufficient Statistics

As Gaussian distribution requires only mean and covariance to be uniquely identified, hence data in classes can be summarized by their sufficient statistics; sample means and covariances.

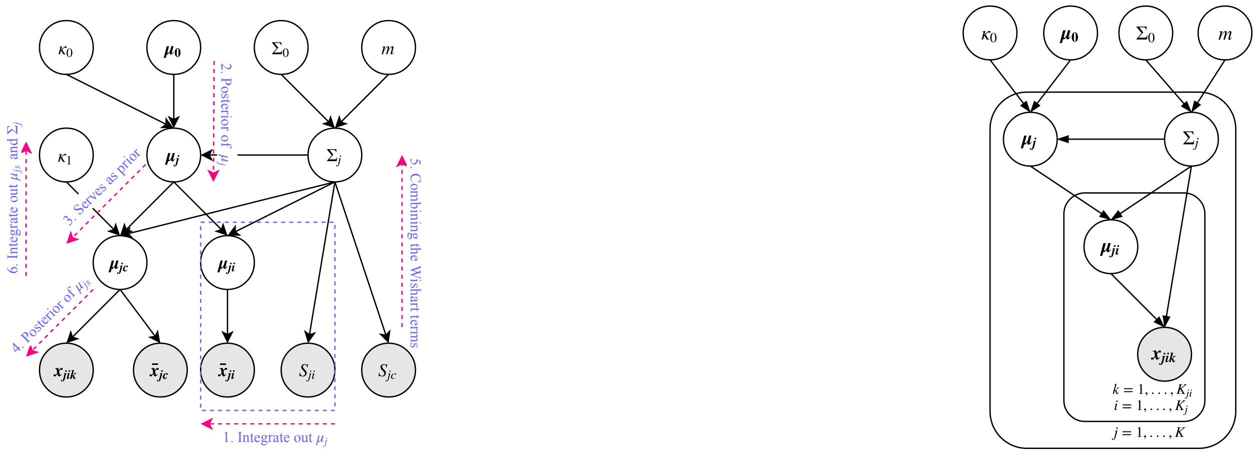


(a) Graphical model

(b) Plate diagram

Figure 3: PPD derivation outline

$$P(\boldsymbol{x_{jik}}|\boldsymbol{\mu_{ji}}, \Sigma_j) \sim N(\boldsymbol{x_{jik}}|\boldsymbol{\mu_{ji}}, \Sigma_j) \tag{1}$$

$$\boldsymbol{\bar{x}_{ji}} = \frac{1}{n_{ji}} \sum_{k:t_k=i} \boldsymbol{x_{jik}} \tag{2}$$

$$\boldsymbol{\bar{x}_{ji}} \sim N(\boldsymbol{\mu_{ji}}, \Sigma_j n_{ji}^{-1}) \tag{3}$$

$$S_{ji} = \frac{1}{n_{ji}} \sum_{k:t_k=i} (\boldsymbol{x_{jik}} - \boldsymbol{\bar{x}_{ji}})(\boldsymbol{x_{jik}} - \boldsymbol{\bar{x}_{ji}})^T \tag{4}$$

$$(n_{ji} - 1)S_{ji} \sim W(\Sigma_j, n_{ji} - 1) \tag{5}$$

(3) follows from eq. (2) and independence assumption given local prior parameters. (5) is a very definition of Wishart distribution as $S_{ji}$ is scatter matrix of class $i$ from local prior $j$.

#### Step 1: Marginal Likelihood

The class sample means $\boldsymbol{\bar{x}_{ji}}'s$ are connected to their local prior ($j$) by integrating out the intermediate class parameter $\boldsymbol{\mu_{ji}}$. Note that all three parameters ($\boldsymbol{\bar{x}_{ji}}, \boldsymbol{\mu_{ji}}, \boldsymbol{\mu_j}$) are Normally distributed and terms depend on local prior covariance ($\Sigma_j$) are treated constant for this derivation.

$$P(\bar{\boldsymbol{x}}_{\boldsymbol{ji}}|\boldsymbol{\mu_j}, \Sigma_j, \kappa_1) = \int P(\bar{\boldsymbol{x}}_{\boldsymbol{ji}}|\boldsymbol{\mu_{ji}}, n_{ji}, \Sigma_j) P(\boldsymbol{\mu_{ji}}|\boldsymbol{\mu_j}, \Sigma_j, \kappa_1) d\boldsymbol{\mu_{ji}} \tag{6}$$

$$= \int N(\bar{\boldsymbol{x}}_{\boldsymbol{ji}}|\boldsymbol{\mu_{ji}}, \Sigma_j n_{ji}^{-1}) N(\boldsymbol{\mu_{ji}}|\boldsymbol{\mu_j}, \Sigma_j \kappa_1^{-1}) \tag{7}$$

$$= \int (2\pi)^{-\frac{d}{2}} |\Sigma_j/n_{ji}|^{-\frac{1}{2}} \exp(-\frac{1}{2}(\bar{\boldsymbol{x}}_{\boldsymbol{ji}} - \boldsymbol{\mu_{ji}})^T (\Sigma_j/n_{ji})^{-1} (\bar{\boldsymbol{x}}_{\boldsymbol{ji}} - \boldsymbol{\mu_{ji}})) \tag{8}$$

$$* (2\pi)^{-\frac{d}{2}} |\Sigma_j/\kappa_1|^{-\frac{1}{2}} \exp(-\frac{1}{2}(\boldsymbol{\mu_{ji}} - \boldsymbol{\mu_j})^T (\Sigma_j/\kappa_1)^{-1} (\boldsymbol{\mu_{ji}} - \boldsymbol{\mu_j})) d\boldsymbol{\mu_{ji}}$$

$$= \int C_1 C_2 exp(-\frac{1}{2}(\boldsymbol{\mu_{ji}} - \frac{\kappa_1 \boldsymbol{\mu_j} + n_{ji} \bar{\boldsymbol{x}}_{\boldsymbol{ji}}}{\kappa_1 + n_{ji}})^T (\frac{\Sigma_j}{n_{ji} + \kappa_1})^{-1} (\boldsymbol{\mu_{ji}} - \frac{\kappa_1 \boldsymbol{\mu_j} + n_{ji} \bar{\boldsymbol{x}}_{\boldsymbol{ji}}}{\kappa_1 + n_{ji}}) + C_3) d\boldsymbol{\mu_{ji}} \tag{9}$$

$$= C_1 C_2 \exp(C_3) \int exp(-\frac{1}{2}(\boldsymbol{\mu_{ji}} - \frac{\kappa_1 \boldsymbol{\mu_j} + n_{ji} \bar{\boldsymbol{x}}_{\boldsymbol{ji}}}{\kappa_1 + n_{ji}})^T (\frac{\Sigma_j}{n_{ji} + \kappa_1})^{-1} (\boldsymbol{\mu_{ji}} - \frac{\kappa_1 \boldsymbol{\mu_j} + n_{ji} \bar{\boldsymbol{x}}_{\boldsymbol{ji}}}{\kappa_1 + n_{ji}}) d\boldsymbol{\mu_{ji}} \tag{10}$$

$$P(\bar{\boldsymbol{x}}_{\boldsymbol{ji}}|\boldsymbol{\mu_j}, \Sigma_j, \kappa_1) = C_1 C_2 \exp(C_3) (2\pi)^{\frac{d}{2}} |\frac{\Sigma_j}{\kappa_1 + n_{ji}}|^{\frac{1}{2}} \tag{11}$$

$$C_1 = (2\pi)^{-\frac{d}{2}} |\Sigma_j/n_{ji}|^{-\frac{1}{2}} \tag{12}$$

$$C_2 = (2\pi)^{-\frac{d}{2}} |\Sigma_j/\kappa_1|^{-\frac{1}{2}} \tag{13}$$

$$C_3 = -\frac{1}{2}(\bar{\boldsymbol{x}}_{\boldsymbol{ji}}{}^T (\Sigma_j/n_{ji})^{-1} \bar{\boldsymbol{x}}_{\boldsymbol{ji}} + \boldsymbol{\mu_j}{}^T (\Sigma_j/\kappa_1)^{-1} \boldsymbol{\mu_j} - \frac{\kappa_1 \boldsymbol{\mu_j} + n_{ji} \bar{\boldsymbol{x}}_{\boldsymbol{ji}}}{\kappa_1 + n_{ji}}^T (\frac{\Sigma_j}{n_{ji} + \kappa_1})^{-1} \frac{\kappa_1 \boldsymbol{\mu_j} + n_{ji} \bar{\boldsymbol{x}}_{\boldsymbol{ji}}}{\kappa_1 + n_{ji}}) \tag{14}$$

$$C_3 = -\frac{1}{2}((\bar{\boldsymbol{x}}_{\boldsymbol{ji}} - \boldsymbol{\mu_j})^T \frac{n_{ji} \kappa_1 \Sigma_j^{-1}}{(n_{ji} + \kappa_1)} (\bar{\boldsymbol{x}}_{\boldsymbol{ji}} - \boldsymbol{\mu_j})) \tag{15}$$

$$P(\bar{\boldsymbol{x}}_{\boldsymbol{ji}}|\boldsymbol{\mu_j}, \Sigma_j, \kappa_1) = (2\pi)^{-\frac{d}{2}} |\frac{\Sigma_j(\kappa_1 + n_{ji})}{n_{ji}\kappa_1}|^{-\frac{1}{2}} \exp(-\frac{1}{2}(\bar{\boldsymbol{x}}_{\boldsymbol{ji}} - \boldsymbol{\mu_j})^T \frac{n_{ji} \kappa_1 \Sigma_j^{-1}}{(n_{ji} + \kappa_1)} (\bar{\boldsymbol{x}}_{\boldsymbol{ji}} - \boldsymbol{\mu_j})) \tag{16}$$

$$P(\bar{\boldsymbol{x}}_{\boldsymbol{ji}}|\boldsymbol{\mu_j}, \Sigma_j, \kappa_1, n_{ji}) = N(\bar{\boldsymbol{x}}_{\boldsymbol{ji}}|\boldsymbol{\mu_j}, \Sigma_j(\frac{1}{n_{ji}} + \frac{1}{\kappa_1})) \tag{17}$$

(6), (7) and (8) follow the model assumption and definition of Normal distribution. (9) is derived by completing the (8) into normal distribution and combining extra elements into constant $C_3$. New term in the (11) comes from evaluation of integral from (10) as the exponential term is in the Gaussian form. Combining the similar terms in (14), we get (15), hence marginal likelihood (16). Hereon, it is trivial to observe that the likelihood is in Gaussian form with mean and covarinace as in (17).

#### Step 2: Posterior of $\boldsymbol{\mu_k}$

We combined the sufficient statistics (means) of classes sharing the same local prior in the posterior distribution of surrogate-class mean $\boldsymbol{\mu_j}$.

$$P(\boldsymbol{\mu_j}|\boldsymbol{\mu_0}, \Sigma_j, \kappa_0, \kappa_1, \{\bar{\boldsymbol{x}}_{\boldsymbol{ji}}\}_{t_i=j}) \propto P(\boldsymbol{\mu_j}|\boldsymbol{\mu_0}, \Sigma_j, \kappa_0) \prod_{i:t_i=j} P(\bar{\boldsymbol{x}}_{\boldsymbol{ji}}|\boldsymbol{\mu_j}, \Sigma_j, \kappa_1) \tag{18}$$

$$P(\boldsymbol{\mu_j}|\boldsymbol{\mu_0}, \Sigma_j, \kappa_0, \kappa_1, \{\bar{\boldsymbol{x}}_{\boldsymbol{ji}}\}_{t_i=j}) \propto exp(-\frac{1}{2}(\boldsymbol{\mu_j} - \bar{\boldsymbol{\mu}}_{\boldsymbol{j}})^T(\sum_{i:t_i=j} \frac{n_{ji}\kappa_1}{(n_{ji}+\kappa_1)} + \kappa_0)\Sigma_j^{-1}(\boldsymbol{\mu_j} - \bar{\boldsymbol{\mu}}_{\boldsymbol{j}})) \tag{19}$$

$$P(\boldsymbol{\mu_j}|\boldsymbol{\mu_0}, \Sigma_j, \kappa_0, \kappa_1, \{\bar{\boldsymbol{x}}_{\boldsymbol{ji}}\}_{t_i=j}) = N(\bar{\boldsymbol{\mu}}_{\boldsymbol{j}}, \bar{\kappa}_j^{-1}\Sigma_j) \tag{20}$$

$$\bar{\boldsymbol{\mu}}_{\boldsymbol{j}} = \frac{\sum_{i:t_i=j} \frac{n_{ji}\kappa_1}{(n_{ji}+\kappa_1)} \bar{\boldsymbol{x}}_{\boldsymbol{ji}} + \kappa_0 \boldsymbol{\mu_0}}{\sum_{i:t_i=j} \frac{n_{ji}\kappa_1}{(n_{ji}+\kappa_1)} + \kappa_0} \tag{21}$$

$$\bar{\kappa}_j = (\sum_{i:t_i=j} \frac{n_{ji}\kappa_1}{(n_{ji}+\kappa_1)} + \kappa_0) \tag{22}$$

Applying Bayes rule, posterior can be proportioned as in (18). As local prior mean and sample mean are Normal distributed (from step 1), we get (19). Completing square procedure used in previous step would give the exact normalization, hence, posterior can be written in a closed form of Gaussian as in (20). The last part can also be verified by observing all exponential terms are quadratic, indeed Gaussian.

#### Step 3: Updated prior of $\boldsymbol{\mu_{jc}}$

As new information is available from classes sharing the same local prior, current class mean ($\boldsymbol{\mu_{jc}}$) can leverage this information by updating its prior. Marginalizing out local prior mean $\boldsymbol{\mu_j}$ would render this information propagation as below,

$$P(\boldsymbol{\mu_{jc}}|\boldsymbol{\mu_0}, \Sigma_j, \kappa_0, \kappa_1, \{\bar{\boldsymbol{x}}_{\boldsymbol{ji}}\}_{t_i=j}) = \int P(\boldsymbol{\mu_{jc}}|\boldsymbol{\mu_j}, \Sigma_j, \kappa_1) P(\boldsymbol{\mu_j}|\bar{\boldsymbol{\mu}}_{\boldsymbol{j}}, \bar{\Sigma}_j, \kappa_0, \kappa_1, \{\bar{\boldsymbol{x}}_{\boldsymbol{ji}}\}_{t_i=j}) d\boldsymbol{\mu_j} \tag{23}$$

$$P(\boldsymbol{\mu_{jc}}|\boldsymbol{\mu_0}, \Sigma_j, \kappa_0, \kappa_1, \{\bar{\boldsymbol{x}}_{\boldsymbol{ji}}\}_{t_i=j}) = \int N(\boldsymbol{\mu_{jc}}|\boldsymbol{\mu_j}, \Sigma_j\kappa_1^{-1}) N(\boldsymbol{\mu_j}|\bar{\boldsymbol{\mu}}_{\boldsymbol{j}}, \bar{\Sigma}_j) d\boldsymbol{\mu_j} \tag{24}$$

$$P(\boldsymbol{\mu_{jc}}|\boldsymbol{\mu_0}, \Sigma_j, \kappa_0, \kappa_1, \{\bar{\boldsymbol{x}}_{\boldsymbol{ji}}\}_{t_i=j}) = N(\boldsymbol{\mu_{jc}}|\bar{\boldsymbol{\mu}}_{\boldsymbol{j}}, \bar{\Sigma}_j + \Sigma_j\kappa_1^{-1}) \tag{25}$$

#### Step 4: Posterior on $\boldsymbol{\mu_{jc}}$

Combining the prior from step 3 with current class sample mean $\bar{\boldsymbol{x}}_{\boldsymbol{jc}}$ from step 1, we derive the posterior for current class mean $\boldsymbol{\mu_{jc}}$. Analogously, applying Bayes rule and observing that both distributions are Normal, we obtain an another Gaussian.

$$P(\boldsymbol{\mu_{jc}}|\boldsymbol{\mu_0}, \Sigma_j, \kappa_0, \kappa_1, \{\bar{\boldsymbol{x}}_{\boldsymbol{ji}}\}_{t_i=j}, \bar{\boldsymbol{x}}_{\boldsymbol{jc}}) \tag{26}$$

$$\propto P(\boldsymbol{\mu_{jc}}|\boldsymbol{\mu_0}, \Sigma_j, \kappa_0, \kappa_1, \{\bar{\boldsymbol{x}}_{\boldsymbol{ji}}\}_{t_i=j}) P(\bar{\boldsymbol{x}}_{\boldsymbol{jc}}|\boldsymbol{\mu_{jc}}, \Sigma_j n_{jc}^{-1}) \tag{27}$$

$$\propto N(\boldsymbol{\mu_{jc}}|\bar{\boldsymbol{\mu}}_{\boldsymbol{j}}, \bar{\Sigma}_j + \Sigma_j \kappa_1^{-1}) N(\bar{\boldsymbol{x}}_{\boldsymbol{jc}}|\boldsymbol{\mu_{jc}}, \Sigma_j n_{jc}^{-1}) \tag{28}$$

$$P(\boldsymbol{\mu_{jc}}|\boldsymbol{\mu_0}, \Sigma_j, \kappa_0, \kappa_1, \{\bar{\boldsymbol{x}}_{\boldsymbol{ji}}\}_{t_i=j}, \bar{\boldsymbol{x}}_{\boldsymbol{jc}}) = N(\frac{n_{jc}\bar{\boldsymbol{x}}_{\boldsymbol{jc}} + \tilde{\kappa}_j \bar{\boldsymbol{\mu}}_{\boldsymbol{j}}}{n_{jc} + \tilde{\kappa}_j}, \Sigma_j(\tilde{\kappa}_j^{-1} + n_{jc}^{-1})) \tag{29}$$

$$\tilde{\kappa}_j^{-1} = \bar{\kappa}_j^{-1} + \kappa_1^{-1} \tag{30}$$

Note that in order to simplify the notation, $\kappa$ terms are collected in $\tilde{\kappa}_j^{-1}$.

Before stepping into the covariance related derivations, we would like to draw your attention to an expression similar to $C_3$ from equation (15) that appears with $\bar{\boldsymbol{\mu}}_{\boldsymbol{j}}$ in the remaining terms while deriving posterior.

$$(\bar{\boldsymbol{x}}_{\boldsymbol{jc}} - \bar{\boldsymbol{\mu}}_{\boldsymbol{j}})^T \frac{n_{jc}\tilde{\kappa}_j}{(n_{jc} + \tilde{\kappa}_j)} \Sigma_j^{-1} (\bar{\boldsymbol{x}}_{\boldsymbol{jc}} - \bar{\boldsymbol{\mu}}_{\boldsymbol{j}}) = \frac{n_{jc}\tilde{\kappa}_j}{n_{jc} + \tilde{\kappa}_j} tr((\Sigma_j^{-1})(\bar{\boldsymbol{x}}_{\boldsymbol{jc}} - \bar{\boldsymbol{\mu}}_{\boldsymbol{j}})(\bar{\boldsymbol{x}}_{\boldsymbol{jc}} - \bar{\boldsymbol{\mu}}_{\boldsymbol{j}})^T)) \tag{31}$$

Note that $\bar{\boldsymbol{x}}_{\boldsymbol{jc}}$ and $\bar{\boldsymbol{\mu}}_{\boldsymbol{j}}$ are observed values, and the equation depends only on $\Sigma_j$. This formula stays in the exponent creating a factor that contributes to Wishart distribution and is denoted as $P(S_\mu|\Sigma_j)$.

#### Step 5: Wishart Terms

As it is observed from plate diagram and generative model in Figure (3), local prior covariance $\Sigma_j$ is shared among its all classes. Even though Wishart terms are independent of mean variables, the residual terms in the posterior calculations create additional Wishart distributions. These distributions and the ones from data scatter matrices are combined in the posterior of $\Sigma_j$ as below,

$$P(\Sigma_j|\{S_{ji}, \bar{\boldsymbol{x}}_{\boldsymbol{ji}}\}_{t_i=j}, S_{jc}, \bar{\boldsymbol{x}}_{\boldsymbol{jc}}) \propto P(\Sigma_j|\Sigma_0, m) P(S_{jc}|\Sigma_j, n_{jc}) P(S_\mu|\Sigma_j) \prod_{i:t_i=j} P(S_{ji}|\Sigma_j, n_{ji}) \tag{32}$$

$$= IW(\Sigma_0 + \sum_{i:t_i=j} S_{ji} + S_{jc} + S_\mu,\ m + \sum_{i:t_i=j} (n_{ji} - 1) + n_{jc}) \tag{33}$$

$$S_\mu = \frac{n_{jc}\tilde{\kappa}_j}{\tilde{\kappa}_j + n_{jc}} (\bar{\boldsymbol{x}}_{\boldsymbol{jc}} - \bar{\boldsymbol{\mu}}_{\boldsymbol{j}})(\bar{\boldsymbol{x}}_{\boldsymbol{jc}} - \bar{\boldsymbol{\mu}}_{\boldsymbol{j}})^T \tag{34}$$

Since the prior on $\Sigma_j$ is Inverse-Wishart and scatter matrices are Wishart distributed, the posterior will be exactly Inverse-Wishart.

#### Step 6: Integration of remaining parameters

As you notice from Step 4 and 5, the posterior distribution is Normal-Inverse-Wishart and from model assumption, data is Gaussian. Hence, the integration in PPD can analytically be derived. Integration with respect to $\boldsymbol{\mu}$ will render another multivariate Normal distribution due to conjugacy, thereafter integration w.r.t $\Sigma$ completes to Inverse-Wishart. Finally, arranging the terms yields the posterior predictive distribution in the form of Student-t.

$$\begin{aligned}
&P(\boldsymbol{x}|\{\bar{\boldsymbol{x}}_{\boldsymbol{ji}}, S_{ji}\}_{t_i=j}, \bar{\boldsymbol{x}}_{\boldsymbol{jc}}, S_{jc}, \boldsymbol{\mu_0}, \kappa_0, \kappa_1) && (35)\\
&= \int\int P(\boldsymbol{x}|\boldsymbol{\mu_{jc}}, \Sigma_j)P(\boldsymbol{\mu_{jc}}, \Sigma_j|\{\bar{\boldsymbol{x}}_{\boldsymbol{ji}}, S_{ji}\}_{t_i=j}, \bar{\boldsymbol{x}}_{\boldsymbol{jc}}, S_{jc}, \boldsymbol{\mu_0}, \kappa_0, \kappa_1)d\boldsymbol{\mu_{jc}}d\Sigma_j\\
&= T(\boldsymbol{x}|\bar{\boldsymbol{\mu}}_{\boldsymbol{jc}}, \bar{\Sigma}_{jc}, \bar{v}_{jc}) && (36)
\end{aligned}$$

$$\begin{aligned}
\bar{\boldsymbol{\mu}}_{\boldsymbol{jc}} &= \frac{n_{jc}\bar{\boldsymbol{x}}_{\boldsymbol{jc}} + \tilde{\kappa}_j\bar{\boldsymbol{\mu}}_{\boldsymbol{j}}}{n_{jc} + \tilde{\kappa}_j}\\
\bar{v}_{jc} &= n_{jc} + \sum_{i:t_i=j}(n_{ji} - 1) + m - d + 1\\
\bar{\Sigma}_{jc} &= \frac{n_{jc} + \tilde{\kappa}_j + 1}{(n_{jc} + \tilde{\kappa}_j)\bar{v}_{jc}}(\Sigma_0 + \sum_{i:t_i=j} S_{ji} + S_{jc} + S_\mu)
\end{aligned}$$

Note that aforementioned derivation of seen class PPD includes local priors, yet in our experiments we simply dropped local priors from Seen class PPD. Hence, the final PPD for seen classes in which local priors are omitted is derived also in the form of a Student-t distribution as following,

$$\begin{aligned}
&P(\boldsymbol{x}|\{\bar{\boldsymbol{x}}_{\boldsymbol{ji}}, S_{ji}\}_{t_i=j}, \bar{\boldsymbol{x}}_{\boldsymbol{jc}}, S_{jc}, \boldsymbol{\mu_0}, \kappa_0, \kappa_1) = T(\boldsymbol{x}|\bar{\boldsymbol{\mu}}_{\boldsymbol{jc}}, \bar{\Sigma}_{jc}, \bar{v}_{jc})\\
&\bar{\boldsymbol{\mu}}_{\boldsymbol{jc}} = \frac{n_{jc}\bar{\boldsymbol{x}}_{\boldsymbol{jc}} + \frac{\kappa_0\kappa_1}{\kappa_0+\kappa_1}\boldsymbol{\mu_0}}{n_{jc} + \frac{\kappa_0\kappa_1}{\kappa_0+\kappa_1}},\ \bar{v}_{jc} = n_{jc} + m - D + 1,\ \bar{\Sigma}_{jc} = \frac{(\Sigma_0 + S_{jc} + S_\mu)(n_{jc} + \frac{\kappa_0\kappa_1}{\kappa_0+\kappa_1} + 1)}{(n_{jc} + \frac{\kappa_0\kappa_1}{\kappa_0+\kappa_1})\bar{v}_{jc}}
\end{aligned} \quad (37)$$

where, $S_\mu$ is defined as in Eq (34). The index $c$ in Equation (37) represents the current seen class, whose PPD is being derived.

Dropping the current class statistics from Eq (36), on the other hand, delivers the PPD for unseen classes as below,

$$\begin{aligned}
&P(\boldsymbol{x}|\{\bar{\boldsymbol{x}}_{\boldsymbol{ji}}, S_{ji}\}_{t_i=j}, \boldsymbol{\mu_0}, \kappa_0, \kappa_1) = T(\boldsymbol{x}|\bar{\boldsymbol{\mu}}_{\boldsymbol{j}}, \bar{\Sigma}_j, \bar{v}_j) && (38)\\
&\bar{v}_j = \sum_{i:t_i=j}(n_{ji} - 1) + m - d + 1, \qquad \bar{\Sigma}_j = \frac{(\tilde{\kappa}_j + 1)}{\tilde{\kappa}_j\bar{v}_j}(\Sigma_0 + \sum_{i:t_i=j} S_{ji})
\end{aligned}$$

where $\bar{\boldsymbol{\mu}}_{\boldsymbol{j}}$ is from (21).

### 3 Training Details

In this section, we provide training details regarding implementation and hyperparameter tuning for CNN model and Bayesian classifier.

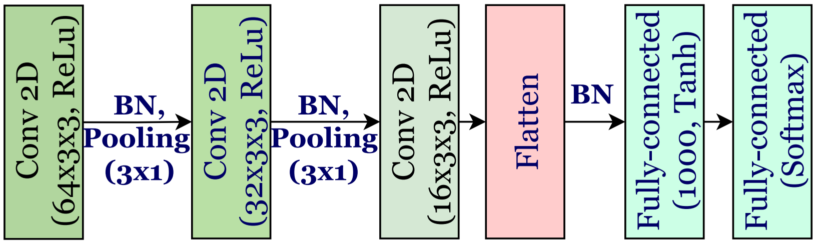


Figure 4: CNN model architecture. BN and Pooling stand for batch normalization and Max-Pooling respectively. Activation functions and layer dimensions are displayed in parenthesis.

#### 3.1 Implementation

Experiments are run on a machine with the following specifications: Dell PC with Windows 10, Intel(R) Core(TM) i9-9900 CPU @ 3.10 GHz, 64 GB RAM, and NVIDIA GeForce RTX 2080 GPU, 8GB RAM.

##### 3.1.1 CNN model

The details of CNN model training are mentioned in the main text, but here we present the detailed model architecture in Figure 4. For the padding of missing bases, we simply used the label *others* which represents missing and ambiguous symbols.

Jupyter notebook together with *Readme* file for learning DNA embeddings is attached to the Supplementary material with the name *DNA_embeddings.zip*. Please refer to the *Readme* file for setting up the *conda* virtual environment and installing required packages in order to run the code.

##### 3.1.2 Bayesian classifier

The model is developed in MATLAB, and any version including 2016 and above should be able to run the model without any problem. The code is attached to the Supplementary material under the same *Bayesian_classifier.zip*. You may find it useful to check the *Readme* file to reproduce reported results.

#### 3.2 Hyperparameter Tuning

##### 3.2.1 CNN Model

We very coarsely tuned the CNN model. After fixing the model architecture, we tuned initial learning rate, batch size and number of epochs between the ranges given below:

- Learning rate: $\{0.1, 0.01, 0.05, 0.001\}$
- Batch size: $\{32, 64\}$
- Number of epochs: $\{5, 10\}$

We also tried *SGD* optimizer but *ADAM* optimizer is superior. With ADAM optimizer CNN model took 52 seconds for training per epoch and less than 5 minutes in total.

##### 3.2.2 Bayesian classifier

Referencing the main text, unconstrained model has 4 parameters to tune: $\{\kappa_0, \kappa_1, m, s\}$. As you may recall from the main text, the hyperparameter $\kappa_0$ adjusts the separation between local prior centers, on the other hand, $\kappa_1$ adjusts the dispersion between class centers inheriting the same local prior. Moreover, $m$ controls the degree of deviation of individual $\Sigma_j$'s from the $E[\Sigma_j]$ and $s$ is the scale constant. As the expected value of $\Sigma_j$ is $\frac{\Sigma_0}{m-d-1}$ where $d$ is the dimension of the data, larger values for $m$ assumes classes with more spherical shapes, whereas smaller values creates resilience to learn more flexible shapes.

Model hyperparameters are coarsely tuned to maximize the Harmonic mean of seen and unseen class accuracies on validation set. The only preprocessing we did was to apply PCA to reduce the dimensionality of the data from 2048 to 500. The range of each parameter and the best quadruplets associated with each model are depicted in the Tables 2 and 3, respectively. Bayesian classifier took 5 seconds to train and 49 seconds for the inference on INSECT dataset. Note that the training time for baseline bioinformatics method is more than 4 hours.

| HP | Range |
|---|---|
| $\kappa_0$ | $\{0.1, 1\}$ |
| $\kappa_1$ | $\{1, 10\}$ |
| $m$ | $\{2d, 5d, 25d, 100d\}$ |
| $s$ | $\{0.5, 1, 5\}$ |

Table 2: Parameter ranges used in hyper-parameter tuning.

| Models | Best quadruplets |
|---|---|
| OSBC-IMG | $\{0.1, 10, 5d, 1\}$ |
| OSBC-DNA | $\{0.1, 10, 25d, 0.5\}$ |
| OSBC-DIL | $\{0.1, 10, 5d, 0.5\}$ |
| OSBC-DIT | $\{0.1, 10, 25d, 0.5\}$ |

Table 3: Best quadruplets from tuning within the order of $\{\kappa_0, \kappa_1, m, s\}$.

### 4 Notes on Model Limitations

We observed that when the DNA barcodes of the species are of a very low quality and have many missing bp's, the embeddings from CNN model are not very informative. These instances, in turn, can lead trivial misclassification cases where misclassified species do not share a common morphology with the test species. The underlying reason for this phenomenon is the following: in the embedding space, some species from training set cluster far apart from their corresponding species belonging to the same genera. These samples, in turn, shift the surrogate-class formation of that genus and distort its distribution. During the inference, the bayesian model fails to classify the samples of undescribed species from that genus to the correct surrogate genus class but misclassifies it to completely different genus.

A more robust embeddings of DNA barcodes towards missing bp's can be obtained via incorporating self-supervised loss into CNN model training. Contrastive learning from computer vision domain can be a good candidate to explore this approach.
